# A broadly generalizable stabilization strategy for sarbecovirus fusion machinery vaccines

**DOI:** 10.1101/2023.12.12.571160

**Authors:** Jimin Lee, Cameron Stewart, Alexandra Schaefer, Elizabeth M. Leaf, Young-Jun Park, Daniel Asarnow, John M. Powers, Catherine Treichel, Davide Corti, Ralph Baric, Neil P. King, David Veesler

## Abstract

Continuous evolution of SARS-CoV-2 alters the antigenicity of the immunodominant spike (S) receptor-binding domain and N-terminal domain, undermining the efficacy of vaccines and monoclonal antibody therapies. To overcome this challenge, we set out to develop a vaccine focusing antibody responses on the highly conserved but metastable S_2_ subunit, which folds as a spring-loaded fusion machinery. Here, we describe a protein design strategy enabling prefusion-stabilization of the SARS-CoV-2 S_2_ subunit and high yield recombinant expression of trimers with native structure and antigenicity. We demonstrate that our design strategy is broadly generalizable to all sarbecoviruses, as exemplified with the SARS-CoV-1 (clade 1a) and PRD-0038 (clade 3) S_2_ fusion machineries. Immunization of mice with a prefusion-stabilized SARS-CoV-2 S_2_ trimer vaccine elicits broadly reactive sarbecovirus antibody responses and neutralizing antibody titers of comparable magnitude against Wuhan-Hu-1 and the immune evasive XBB.1.5 variant. Vaccinated mice were protected from weight loss and disease upon challenge with SARS-CoV-2 XBB.1.5, providing proof-of-principle for fusion machinery sarbecovirus vaccines motivating future development.

## Introduction

Several COVID-19 vaccines have been authorized worldwide to induce antibody responses targeting the SARS-CoV-2 spike (S) glycoprotein. These vaccines enabled safe and effective protection against infection with the Wuhan-Hu-1 (Wu) isolate, which swept the globe at the beginning of the COVID-19 pandemic. However, continued viral evolution led to the emergence of SARS-CoV-2 variants with distinct antigenic properties, relative to previous isolates, eroding neutralizing antibody responses^1^. As a result, breakthrough infections have become common although^2–6^ vaccinated individuals remain protected from severe diseases^7–12^. Furthermore, the neutralizing activity of monoclonal antibody therapies has been compromised by these antigenic changes, resulting in withdrawal of their regulatory authorization.

The SARS-CoV-2 S glycoprotein receptor-binding domain (RBD) is targeted by a vast diversity of antibodies and RBD-directed antibodies account for most of the plasma neutralizing activity against infection/vaccine-matched and mismatched viruses^13–15^. Conversely, the S N-terminal domain (NTD) is mostly targeted by variant-specific neutralizing antibodies^3,16,17^. The SARS-CoV-2 S_2_ subunit (fusion machinery) is much more conserved than the S_1_ subunit (comprising the RBD and NTD), and harbors several antigenic sites targeted by broadly reactive monoclonal antibodies, including the stem helix^18–20^, the fusion peptide^21–23^ and the trimer apex^24^. Although some of these antibodies have neutralizing activity against a wide range of variants and distantly related coronaviruses, and protect small animals against viral challenge, their potency is limited compared to best-in-class RBD-directed antibodies^25–27^. Furthermore, fusion machinery-directed antibodies are rare in the plasma and memory B cells of previously infected and/or vaccinated subjects and have limited contribution to neutralization mediated by polyclonal antibodies^14,18^. Therefore, vaccines enabling to overcome this challenge through elicitation of high titers of neutralizing antibodies targeting the conserved S_2_ subunit bear the promise to limit the need for vaccine update.

Here, we set out to design a prefusion-stabilized SARS-CoV-2 S_2_ subunit vaccine enabling robust elicitation of antibodies targeting conserved fusion machinery antigenic sites to limit the impact of viral evolution on immune responses. Coronavirus S glycoproteins fold as spring-loaded trimers transitioning from the prefusion to the postfusion state, upon receptor engagement and proteolytic activation to promote membrane fusion and viral entry^28–30^. This metastability, however, constitutes a challenge for producing the S_2_ subunit (fusion machinery) in the prefusion state in the absence of the S_1_ subunit. We identified a set of mutations allowing high-yield recombinant production of fusion machinery trimers with native prefusion architecture and antigenicity that remained stable in various storage conditions, as revealed through structural and serological studies. We further show that the prefusion-stabilization strategy designed is broadly generalizable to sarbecovirus S_2_ subunits and successfully ported the identified mutations to the SARS-CoV-1 and PRD-0038 fusion machineries to generate vaccine candidates with optimal biophysical properties. Immunization of mice with a designed SARS-CoV-2 fusion machinery trimer vaccine elicited broadly reactive sarbecovirus antibodies and neutralizing antibody titers of comparable magnitude against the Wu and the immune evasive XBB.1.5 variant. Vaccinated mice were protected from weight loss and disease upon challenge with SARS-CoV-2 XBB.1.5 motivating further development of this vaccine.

## Results

### Design of prefusion-stabilized SARS-CoV-2 S_2_ subunit vaccines

We previously designed a fusion machinery (S_2_ subunit) antigen^14^ stabilized through introduction of 4 out of 6 HexaPro mutations (A892P, A899P, A942P and V987P)^31^, the VFLIP inter-protomer disulfide (Y707C-T883C)^32^ and an intra-protomer disulfide (F970C-G999C) (**Figure 1A**). The resulting construct (designated C-44) harbored protomers with a prefusion tertiary structure but a quaternary structure in which the viral membrane distal region (apex) was splayed open compared to prefusion SARS-CoV-2 S^14^ (**Figure 1B**). To design a prefusion S_2_ subunit trimer with a native quaternary structure, we selected mutations from a deep-mutational scanning dataset of cell-surface expressed S using a library spanning residues 883 to 1034^33^. Mutations were ranked according to their expression/fusion score ratio, to favor prefusion-stabilizing amino acid substitutions, and visually inspected to lead to a final selection of ten mutations (**Figure 1A, Supplementary Table 1**). We first evaluated the effect of three individual mutations introduced in the C-44 background, namely T961F, D994E, and Q1005R, which were among the highest ranking based on their expression/fusion score ratio. We recombinantly produced each of the three constructs (designated E-31, E-32 and E-33 for the T961F, D994E and Q1005R mutations, respectively) in human cells and characterized their monodispersity by size-exclusion chromatography (SEC). Residue 961 is part of the heptad repeat 1 (HR1) motif whereas residues 994 and 1005 are located in the central helix (CH) ^28,29,34^. All three constructs eluted mainly as a monodisperse species, yielding 40-60 mg of purified protein per liter of Expi293 cells (**Figure 1C and Figure 1D)**. Single-particle electron microscopy (EM) characterization of each negatively stained glycoprotein revealed that the design harboring the T961F mutation (E-31) formed prefusion closed S_2_ trimers whereas the other two constructs, harboring either D994E (E-32) or Q1005R (E-33), adopted several conformations, including the one with a splayed-open apex (**Supplementary Figure 1**), as previously observed for C-44^14^. Furthermore, we observed a higher tendency to aggregate for E-32 and E-33 by SEC and EM (**Figure 1C and Supplementary Figure 1**). To unveil the E-31 architecture and confirm the effect of T961F mutation, we determined a cryoEM structure of this antigen at 2.7 Å resolution (**Figure 1E, Supplementary Figure 2 and Supplementary Table 2)**. Our structure shows that E-31 folds with virtually identical tertiary and quaternary structures to the S_2_ subunit from the prefusion S ectodomain trimer (PDB 6VXX^34^) with which the protomers can be superimposed with r.m.s.d values of 0.7 Å. The engineered T961F substitution, which maps to the distal half of the fusion machinery apex, reinforces interactions between HR1 and the central helix of one protomer and the upstream helix of a neighboring protomer (cavity filling), likely explaining the compact, closed trimer conformation observed (**Supplementary Figure 3A)**. Although we could detect a small fraction of E-31 trimers with an open apex conformation in the cryoEM dataset, our findings demonstrate that the T961F mutation alone is sufficient to close the fusion machinery apex in a prefusion conformation (**Figure 1E, Supplementary Figure 2**).

**Figure 1.**
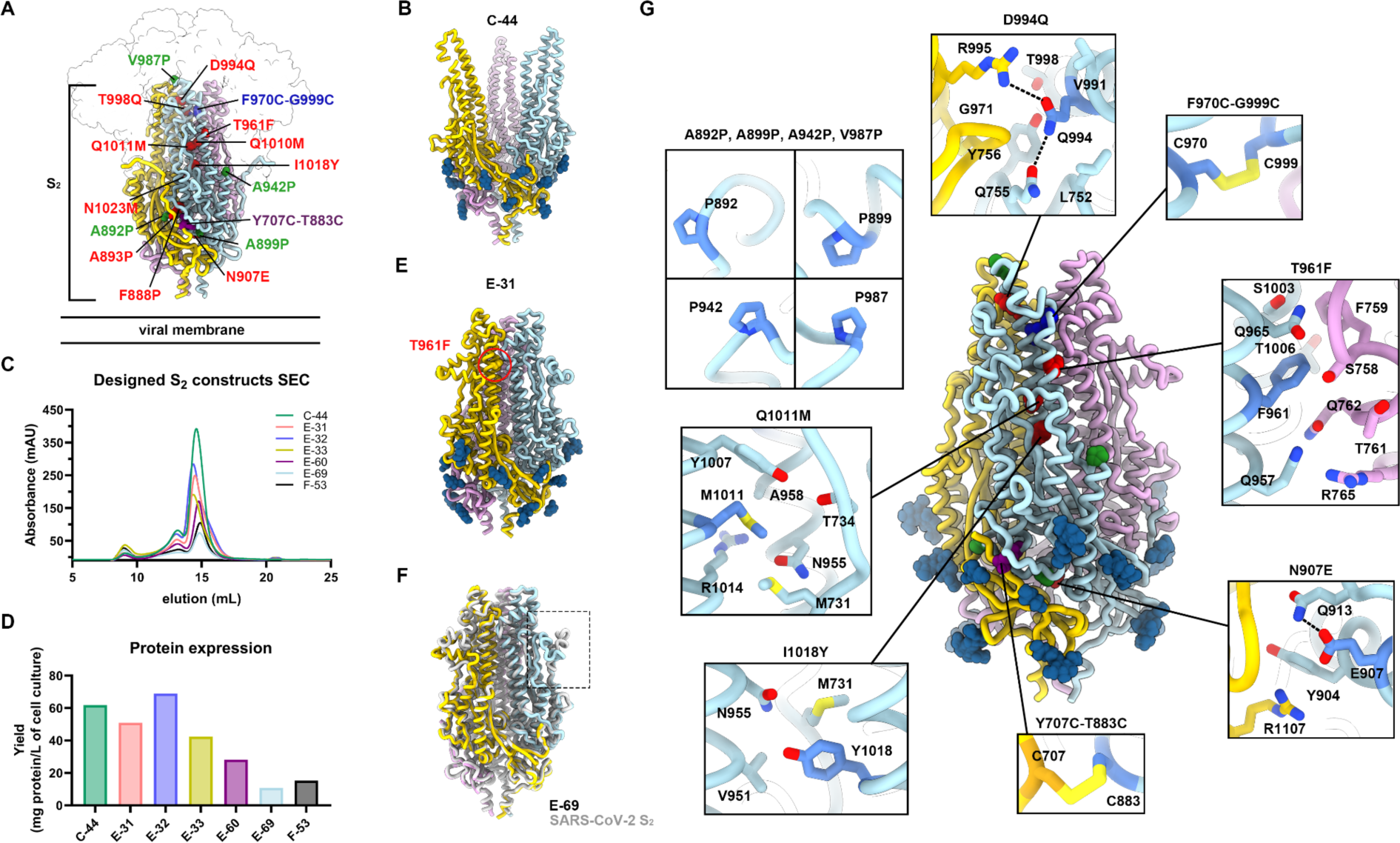
Design of prefusion-stabilized SARS-CoV-2 fusion machinery (S_2_ subunit) vaccines. **A,** Ribbon diagram of prefusion SARS-CoV-2 S highlighting all positions that were mutated to attempt to stabilize the metastable fusion machinery (S_2_ subunit) in the prefusion conformation. Mutations are shown in blue (intra-protomer disulfide bond), purple (VFLIP inter-protomer disulfide bond^32^), red (ten mutations selected based on expression/fusion score of a deep-mutational scan^33^), and green (subset of proline mutations selected from HexaPro^31^). The S_1_ subunit is shown as a transparent surface and glycans are omitted for clarity. **B,** Ribbon diagram of the C-44 cryoEM structure previously determined^14^ (PDB 8DYA). **C,** Size-exclusion chromatograms of the designed S_2_ constructs. **D,** Purification yields of the designed S_2_ constructs after size-exclusion chromatography. **E,** Ribbon diagram of the E-31 cryoEM structure. The position of the T961F mutation is circled red in one protomer. **F,** Superimposition of the S_2_ subunits from the E-69 cryoEM structure and prefusion SARS-CoV-2 S^34^ (gray, PDB 6XR8, residues 704-1146). The box denotes a local structural deviation. Glycans are omitted for clarity. **G,** Ribbon diagram of the E-69 cryoEM structure. Insets: zoomed-in views of the mutations introduced with mutated residues rendered with darker blue/orange shades. Selected electrostatic interactions are shown as dashed lines. The three protomers of each trimer are colored blue, pink and gold throughout the figure.

To further improve the conformational homogeneity of our vaccine candidate, we designed two additional constructs comprising either 4 or 9 additional residue substitutions added to the E-31 trimer (**Supplementary Table 1**). Both constructs were recombinantly produced to characterize their expression, stability and structural properties (**Figure 1C and Figure 1D**). The E-60 construct harbors all 9 additional mutations we selected on the E-31 scaffold. Although recombinant production of this protein construct led to high expression levels, its cryoEM structure at 3.5 Å resolution (**Supplementary Figure 4, Supplementary Table 2**) revealed that the introduction of several proline mutations in the region comprising residues 879-897 distorted an α-helix and following loop relative to its native structure **(Supplementary Figure 3B)**. Ruling out conformation distorting-mutations and a few additional mutations based on visual inspection, we designed a new construct which we named E-69 and harbors the N907E, D994Q, Q1011M and I1018Y in the E-31 background. To investigate its atomic-level organization, we determined a cryoEM structure of E-69 at 3Å resolution and observed clear cryoEM density defining the side chains of all amino acid mutations introduced and confirming folding as a prefusion closed S_2_ subunit trimer, without particle images corresponding to trimers with an open apex (**Figure 1F and Supplementary Figure 5**). The overall E-69 architecture is nearly identical to that of the S_2_ subunit trimer from the prefusion S ectodomain structure (PDB 6VXX^34^) (**Figure 1F**). Only one region noticeably deviated from prefusion S in terms of local structure: a loop downstream of the fusion peptide (residues 827-856) (**Figure 1F and Supplementary Figure 3C**). In the intact SARS-CoV-2 S trimer, this region makes contact with domains C and D of a neighboring protomer which is absent in our design, likely allowing the region to adopt an altered local conformation (**Supplementary Figure 3C**).The N907E substitution places the newly introduced side chain carboxylate close to the Q913 side chain amide with which it interacts electrostatically (**Figure 1G**). The D994Q side chain amide forms an intra-protomer hydrogen bond with the Q755 side chain amide and an inter-protomer electrostatic interaction with the R995 guanidinium, thereby strengthening interactions at the distal part of the apex (**Figure 1G**). The Q1011M mutation is likely stabilizing through reinforcement of local hydrophobic packing with the nearby M731 and Y1007 residues (**Figure 1G**). The I1018Y mutation fills a cavity within each protomer through local interactions involving the CH, HR1 and upstream helix region (**Figure 1G**).

As our E-69 structure did not resolve the flexible N-terminal region, which connects the S_1_ and S_2_ subunits in the context of the S trimer, we designed a new construct lacking 15 residues at the N-terminus of E-69 and named it F-53 (**Figure 1C, 1D, Supplementary Table 1**). F-53 expressed better than E-69 and retained identical antigenicity, and thus served as a template for subsequent S_2_ antigen design (**Supplementary Figure 6**).

### A stable prefusion-stabilized SARS-CoV-2 fusion machinery antigen

Storage and shipping conditions are important considerations for vaccine design and manufacturing as they can impact the cost of goods, ease of distribution and in turn global access. To evaluate the stability of the prefusion-stabilized SARS-CoV-2 E-69 design, we investigated retention of antigenicity after storage in various conditions by ELISA using a panel of monoclonal antibodies targeting the S_2_ subunit (**Figure 2A, 2B**). The stem helix-targeting S2P6 antibody^18^, the fusion peptide-directed 76E1 antibody^22^, and fusion machinery apex recognizing RAY53 bound to E-69, as was the case with SARS-CoV-2 S (**Figure 2A, 2B, and 2C**). We found that E-69 retained unaltered antigenicity for at least two weeks at room temperature and at 4°C, and could be frozen and thawed without affecting its antigenicity (**Figure 2C and Supplementary Figure 7**). To assess the biophysical stability of E-69, we analyzed the purified constructs at various time points using negative staining EM. 2D class averages of E-69 stored in diverse conditions at various time points showed that E-69 retained its prefusion conformation for at least two weeks both at room temperature and at 4°C and that it could be frozen and thawed without altering its structure (**Figure 2D-2I**). Although we detected a minor population of trimers with an open apex upon storage at low temperatures (**Figure 2G-2I**), the 2D class averages suggested that the magnitude of these structural changes was small compared to that observed with the C-44, E-32 or E-33 designed constructs (**Figure 1B, Supplementary Figure 1C, 1D**). Furthermore, we did not detect any 2D classes corresponding to open trimers in our E-69 cryoEM dataset **(Supplementary Figure 5)**. These data suggest that our E-69 design is stable, highlighting the robustness of our prefusion-stabilization strategy, and endowed with optimal biophysical properties for a vaccine candidate.

**Figure 2.**
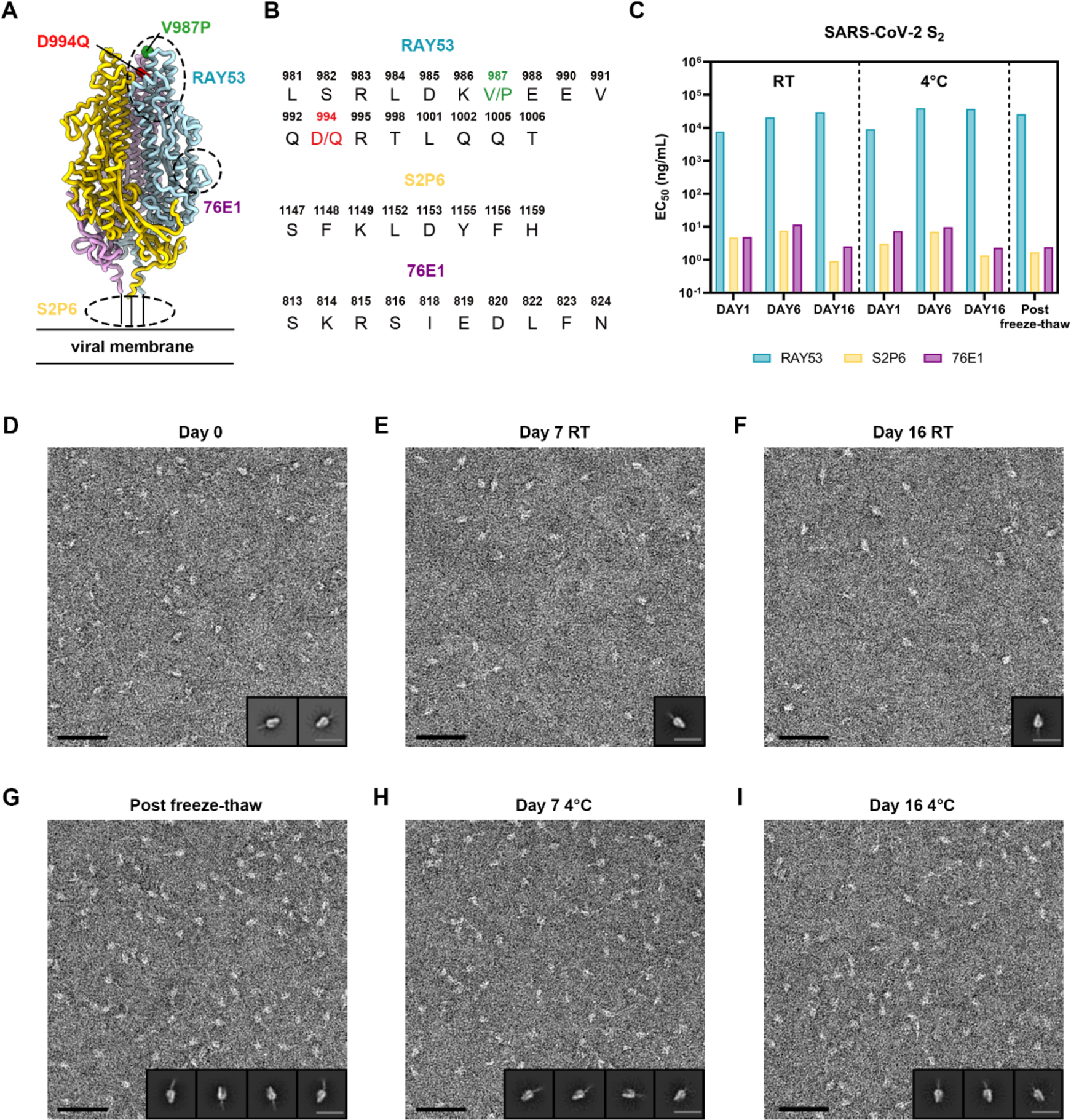
A stable prefusion-stabilized SARS-CoV-2 fusion machinery (S_2_ subunit) vaccine candidate. **A,** Ribbon diagram of the E-69 prefusion-stabilized SARS-CoV-2 fusion machinery antigen highlighting the regions containing the epitopes recognized by S_2_ subunit-targeting monoclonal antibodies using dashed lines. **B,** Amino acid sequences of the RAY53, S2P6 and 76E1 epitopes. The V987P and D994Q E-69 mutations are respectively shown in green and red in panels A-B as they are located within the RAY53 epitope that has been previously reported^24^. **C,** Evaluation of retention of antigenicity for the E-69 antigen in various storage conditions using binding of the S2P6, 76E1 and RAY53 monoclonal antibodies analyzed by ELISA. **D-H,** Evaluation of retention of the native prefusion conformation for the E-69 antigen in various storage conditions analyzed by EM of negatively stained samples. Insets: 2D class averages. The scale bars represent 50 nm (micrograph) and 200 Å (2D class averages).

### A broadly generalizable prefusion-stabilization strategy for sarbecovirus fusion machinery immunogens

To assess the broad applicability of our S_2_ subunit prefusion-stabilization strategy, we examined the local structural environment of the residues mutated in E-69 and compared it with the corresponding regions of interest in SARS-CoV-1 S_2_ (clade 1a) and PRD-0038 S_2_ (clade 3)^35^. The overall architecture of the SARS-CoV-1 S_2_ and PRD-0038 S_2_ trimers is similar to that of SARS-CoV-2 S_2_ with which they can be superimposed with r.m.s.d. values of 1.1 and 1.3 Å, and share 91% and 88% amino acid sequence identity, respectively. Based on the observed conservation of the local structural environment, we hypothesized that the E-69 mutations should be portable to SARS-CoV-1 S_2_ and to PRD-0038 S_2_ and designed the corresponding constructs (**Figure 3A-3I**). We additionally truncated the N-terminal region of the prefusion-stabilized SARS-CoV-1 S_2_ and PRD-0038 S_2_ constructs due to the enhanced expression of SARS-CoV-2 F-53 relative to E-69 (**Figure 1C-D**). Finally, we introduced the V940Q substitution to the PRD-0038 S_2_ construct, which is the position equivalent to SARS-CoV-2 Q957 and SARS-CoV-1 Q939, to allow electrostatic interaction with R748 (similar to that observed with SARS-CoV-2 R765 and SARS-CoV-1 R747 from a neighboring protomer) (**Figure 3D**).

**Figure 3.**
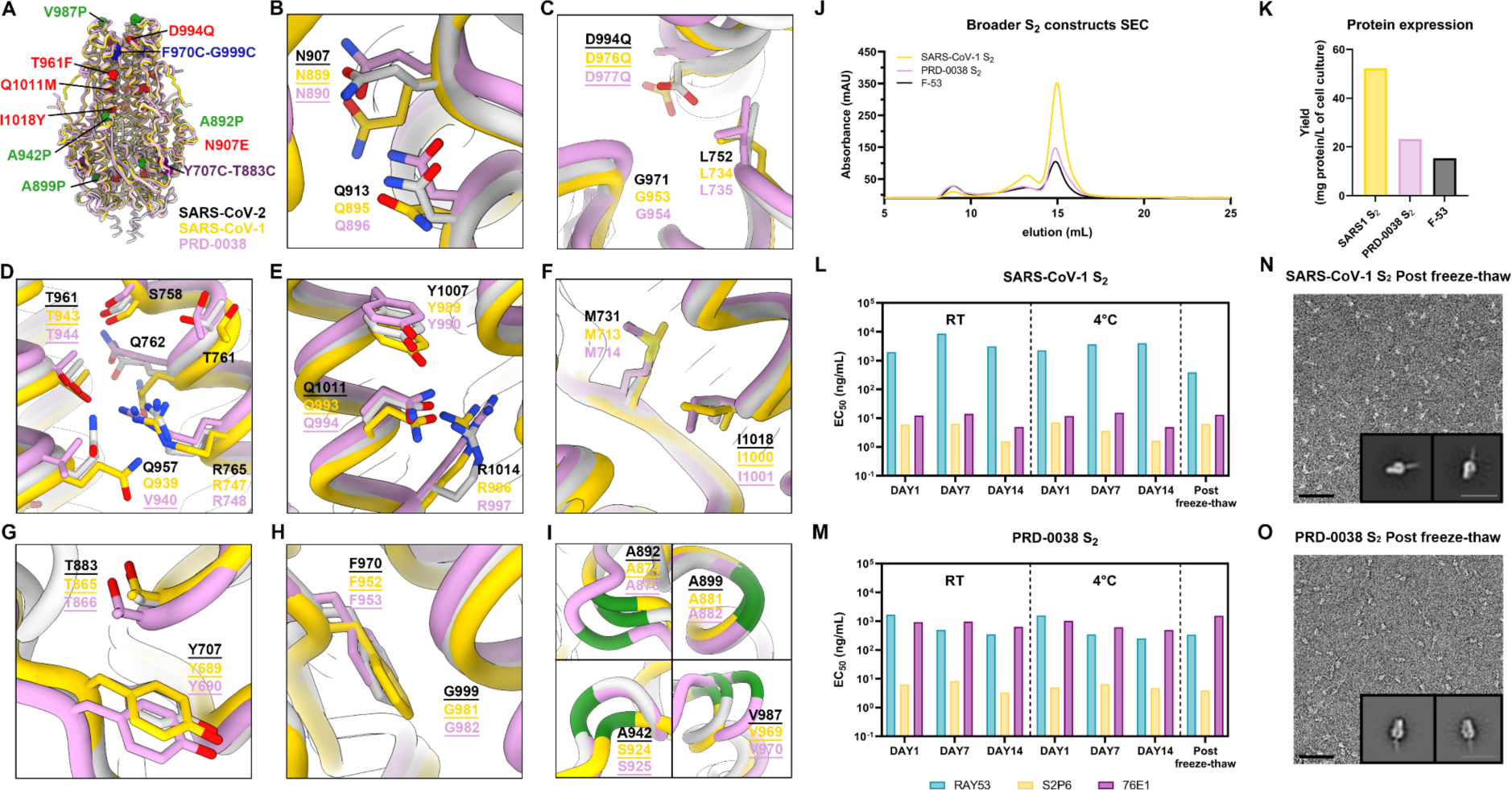
A broadly generalizable prefusion-stabilization strategy for sarbecovirus fusion machinery (S_2_ subunit) antigens. **A,** Ribbon diagrams of superimposed S_2_ subunits of the prefusion SARS-CoV-2 S (PDB 6VXX^34^), SARS-CoV-1 S (PDB 5X58^37^) and PRD-0038 S (PDB 8U29^36^) structures. Prefusion-stabilizing mutations are shown in blue (intra-protomer disulfide bond), purple (VFLIP inter-protomer disulfide bond^32^), red (mutations ported from E-69), and green (subset of proline mutations selected from HexaPro^31^). **B-I,** Zoomed-in views of superimposed S_2_ subunits of the prefusion SARS-CoV-2, SARS-CoV-1 and PRD-0038 S structures highlighting the local structural conservation of residues mutated in SARS-CoV-2 E-69/F-53. Mutated residues in our designed constructs are underlined. SARS-CoV-2, SARS-CoV-1, and PRD-0038 S are respectively shown in light gray, gold, and pink in panels (A-I). **J,** Size-exclusion chromatograms of the designed SARS-CoV-1 and PRD-0038 S_2_ constructs, as compared to SARS-CoV-2 F53. **K,** Purification yields of the designed SARS-CoV-1 and PRD-0038 S_2_ constructs. The yield for the best SARS-CoV-2 S_2_ construct (F53) is included for comparison. **L,M,** Evaluation of retention of antigenicity for the SARS-CoV-1 (L) and PRD-0038 (M) S_2_ antigens in various storage conditions using binding of the S2P6, 76E1 and RAY53 monoclonal antibodies analyzed by ELISA. **N,O,** Evaluation of retention of the native prefusion conformation of the negatively stained SARS-CoV-1 (N) and PRD-0038 (O) S_2_ trimers after freeze/thawing. Insets: 2D class averages showing compact prefusion S_2_ trimers. The scale bar represents 50 nm (micrographs) and 200 Å (2D class averages).

Recombinant production and purification of the designed SARS-CoV-1 and PRD-0038 S_2_ subunit constructs led to even greater yields than that of SARS-CoV-2 F-53, reaching 50 mg of purified SARS-CoV-1 S_2_ trimer per liter of Expi293 cells. (**Figure 3J and 3K**). These two trimers were stable in a variety of storage conditions and retained unaltered reactivity with fusion machinery-directed monoclonal antibodies for at least two weeks at room temperature and at 4°C and could be frozen and thawed without affecting their antigenicity (**Figure 3L and 3M**). Although the RAY53 and S2P6 antibodies each cross-reacted with comparable efficiency to the SARS-CoV-1 S_2_ trimer and to the PRD-0038 S_2_ trimer, 76E1 bound efficiently to SARS-CoV-1 S_2_ but much more weakly to PRD-0038 S_2_, possibly as a result of the F823_SARS-CoV-2_Y806_PRD-0038_ epitope mutation^36^ (**Figure 3L and 3M**). Single particle EM analysis of negatively stained SARS-CoV-1 S_2_ and PRD-0038 S_2_ showed that they adopt the designed closed prefusion architecture (**Figure 3N and 3O**). Collectively, these data indicate that our S_2_ prefusion-stabilization strategy is broadly applicable across different sarbecovirus clades and promotes retention of native structure and antigenicity over time under various storage conditions.

### A prefusion-stabilized SARS-CoV-2 fusion machinery vaccine elicits broadly reactive antibody responses

To evaluate the immunogenicity of our lead prefusion SARS-CoV-2 S_2_ designed vaccine candidate, we immunized twelve BALB/c mice with four 5 µg doses of E-69 on weeks 0, 3, 10 and 17 ands twelve BALB/c mice with two 1 µg doses of prefusion SARS-CoV-2 2P S on weeks 0 and 3 followed by two 5 µg doses of E-69 on weeks 10 and 17. The latter group aims to recapitulate the pre-existing immunity found in humans due to prior exposures through vaccination and/or infection. For benchmarking, we also immunized twelve BALB/c mice with four 1 µg doses of prefusion SARS-CoV-2 2P S on weeks 0, 3, 10 and 17 **(Figure 4A)**. All immunogens were adjuvanted with Addavax using a 1:1 (v/v) ratio.

**Figure 4.**
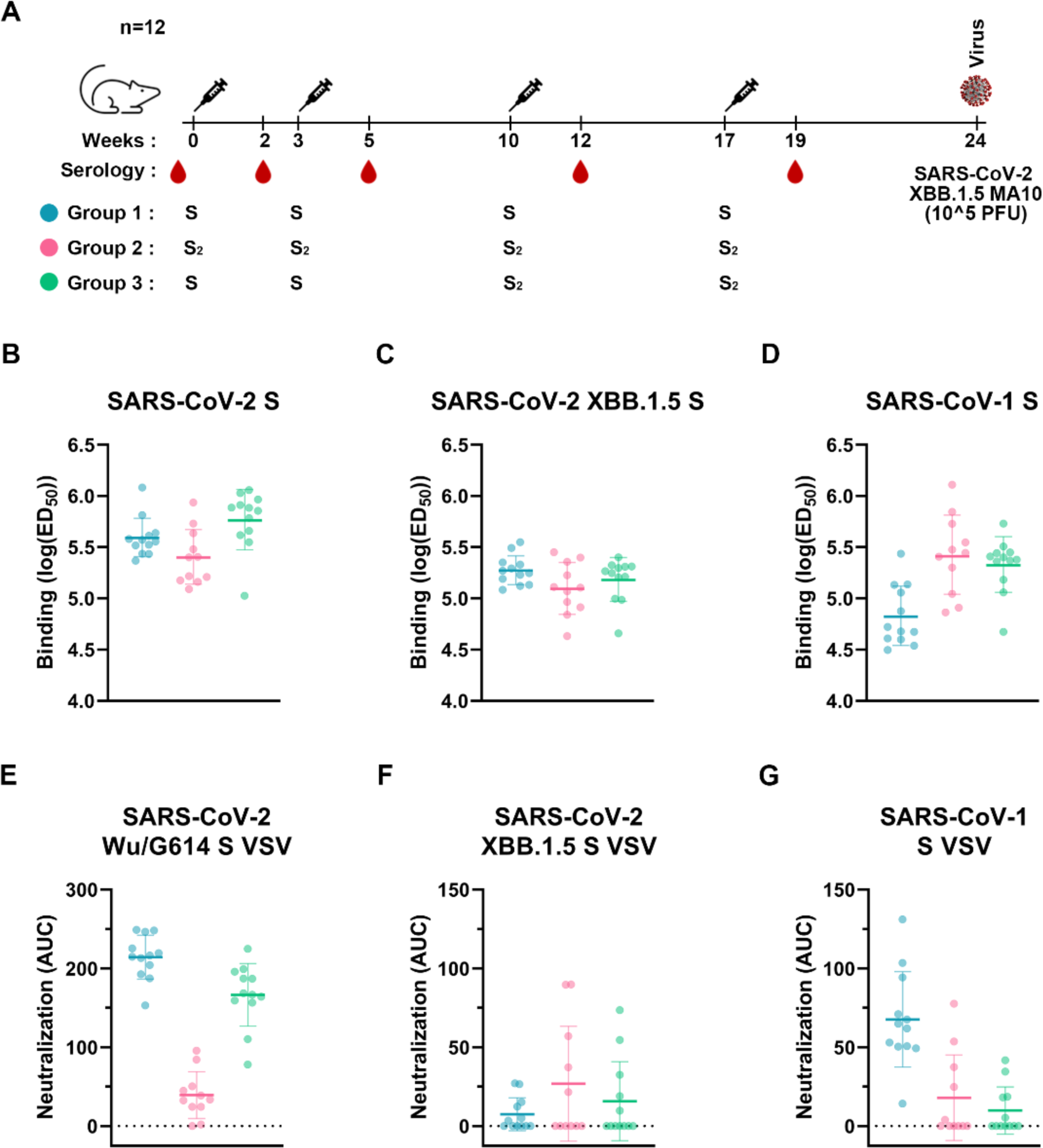
A prefusion-stabilized SARS-CoV-2 fusion machinery (S_2_ subunit) vaccine elicits broadly reactive antibody responses. **A,** Vaccination schedule and study design. **B-D,** Analysis of antibody binding titers against SARS-CoV-2 Hexapro S (B), XBB.1.5 Hexapro S (C), and SARS-CoV-1 Hexapro S (D) analyzed by ELISAs using sera obtained two weeks post dose 4. Geometric means and geometric SD are shown as bars. Representative graphs from two biological replicates are shown. **E-G,** Analysis of neutralizing antibody titers against SARS-CoV-2 Wu/G614 (E), XBB.1.5 (F), and SARS-CoV-1 (G) S VSV pseudoviruses using sera obtained two weeks post dose 4 expressed as area under the curve (AUC). Mean values and SD shown as bars. Baselines at 0 AUC represent no neutralization. Representative graphs from three biological replicates are shown.

Binding antibody titers were analyzed by ELISA using sera obtained two weeks post dose 4 (week 19) against SARS-CoV-2 Wu HexaPro S, SARS-CoV-2 XBB.1.5 HexaPro S, and SARS-CoV-1 HexaPro S **(Figure 4B-D and Supplementary Figure 9)**. We observed similar prefusion SARS-CoV-2 Wu S and XBB.1.5 S antibody binding titers upon immunization with E-69 (GMT: 5.4/5.1), SARS-CoV-2 2P S (GMT: 5.6/5.3) or SARS-CoV-2 2P S followed by E-69 (GMT: 5.8/5.2). However, we observed higher SARS-CoV-1 S binding titers with the E-69 (GMT: 5.4) or SARS-CoV-2 2P S followed by E-69 (GMT: 5.3) vaccination regimens as compared to SARS-CoV-2 2P S immunization (GMT: 4.8). These results suggest that E-69 vaccination elicits comparable serum antibody binding titers after four doses to the widely used prefusion S 2P trimer against SARS-CoV-2 variants but higher titers against SARS-CoV-1 through focusing of antibody responses on the fusion machinery in these experimental conditions.

We subsequently evaluated vaccine-elicited neutralizing activity using sera obtained two weeks post dose 4 (week 19) and vesicular stomatitis virus (VSV) particles pseudotyped with SARS-CoV-2 Wu/G614, XBB.1.5 S or SARS-CoV-1 S (**Figure 4E-4G and Supplementary Figure 10**). SARS-CoV-2 2P S or SARS-CoV-2 2P S followed by E-69 vaccination elicited potent neutralizing antibody titers against the vaccine-matched SARS-CoV-2 Wu/G614 with respective AUCs of 214 and 166 whereas E-69 elicited more modest neutralizing activity with AUC of 40. SARS-CoV-1 S VSV neutralization was highest upon vaccination with SARS-CoV-2 2P S (AUCs of 68) and comparable for the groups vaccinated with E-69 (AUC of 18) or SARS-CoV-2 2P S followed by E-69 (AUC of 10). Neutralization of XBB.1.5 VSV S, however, was highest for E-69-vaccinated mice (AUC of 27) as compared to vaccination with SARS-CoV-2 2P S (AUC of 7) or vaccination with SARS-CoV-2 2P S followed by E-69 (AUC of 16).

### A prefusion-stabilized SARS-CoV-2 fusion machinery vaccine protects against the SARS-CoV-2 XBB.1.5 variant

To study the in vivo protective efficacy of the designed fusion machinery SARS-CoV-2 vaccine against an immune evasive SARS-CoV-2 variant, we intranasally inoculated each animal in the three aforementioned vaccinated groups of mice with 10^5^ PFU of XBB.1.5 MA10 **(Figure 4A)**. We also challenged 10 unvaccinated mice as a control group. Weight loss was followed for 4 days post infection **(Figure 5A)** whereas replicating viral titers in the nasal turbinates and lung as well as lung pathology were assessed at 2 and 4 days post challenge **(Figure 5B-D)**. Although none of the immunogens evaluated protected mice from infection, likely due to the systemic delivery route^10,38–40^, all vaccinated mice were protected against weight loss throughout the duration of the experiment, as compared to unvaccinated mice **(Figure 5A).** Furthermore, lung viral load and lung pathology were comparable for all three vaccinated groups and markedly reduced compared to unvaccinated animals. These data indicate that vaccination with our lead prefusion-stabilized SARS-CoV-2 S_2_ subunit (fusion machinery) elicited protection against the highly immune evasive SARS-CoV-2 XBB.1.5 variant in this stringent challenge model **(Figure 5B, 5C, and 5D)**.

**Figure 5.**
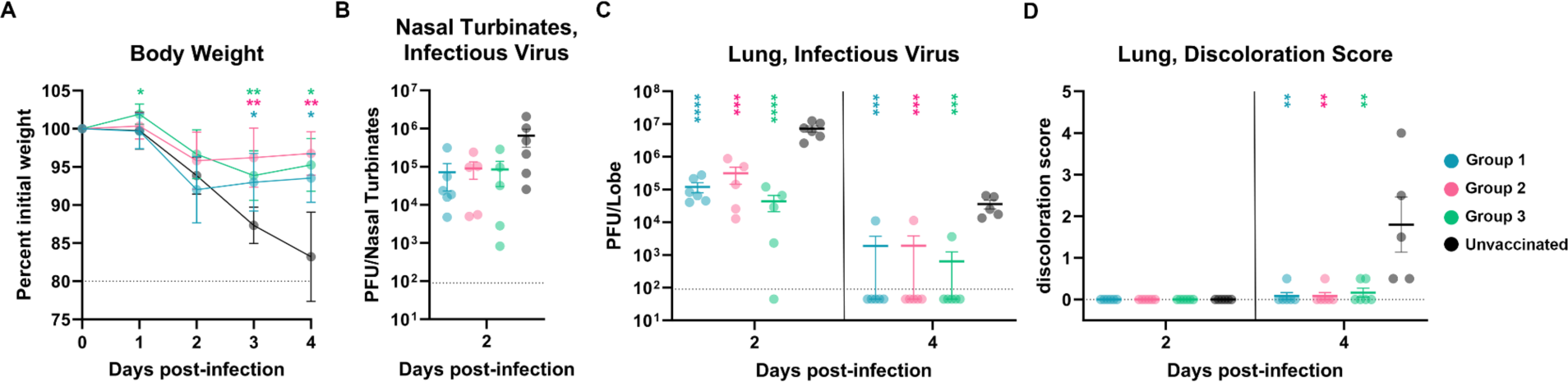
A prefusion-stabilized SARS-CoV-2 fusion machinery (S_2_ subunit) vaccine protects mice against SARS-CoV-2 XBB.1.5-induced disease. **A,** Weight loss followed 4 days post viral challenge with SARS-CoV-2 XBB.1.5 MA10. Control group represents the mice group without vaccination. Comparison of percent weight loss with the control group and each of the vaccinated groups was done using two-way analysis of variance (ANOVA) after Dunnett’s multiple comparison test. *P < 0.05; **P < 0.01; ***P < 0.001; ****P < 0.0001. **B-C,** Quantification of replicating viral titers in the nasal turbinates (B) and lungs (C) of challenged animals at 2 and 4 days post infection, respectively. Mean values and SEM shown as bars. Statistical significance compared with the control group and each of the vaccinated groups was reported using one-way ANOVA after Tukey’s multiple comparison test. *P < 0.05; **P < 0.01; ***P < 0.001; ****P < 0.0001. Limit of detection shown as dashed line. **D,** Lung discoloration score at 2 and 4 days post infection. Mean values and SEM shown as bars. Statistical significance compared with the control group and each of the vaccinated groups was reported using one-way ANOVA after Dunnett’s multiple comparisons test. *P < 0.05; **P < 0.01; ***P < 0.001; ****P < 0.0001.

## Discussion

Emergence of immune evasive SARS-CoV-2 variants erodes the effectiveness of COVID-19 vaccines, which led to the roll out of two updated boosters in 2022^41,42^ and 2023^43^. For the foreseeable future, it is likely that COVID-19 vaccines will require yearly reformulation based on the anticipated prevalence of circulating variants, similarly to influenza virus vaccines. Next-generation vaccines that are more resilient to viral evolution and antigenic changes bear the promise of reducing the need for or the frequency of vaccine updates which would also help with large scale adoption by the public. The sarbecovirus S_2_ subunit prefusion-stabilization strategy presented here represents a key step in this direction due to the much higher conservation of the S_2_ subunit relative to the S_1_ subunit among SARS-CoV-2 variants and other sarbecoviruses^11,34,44^. Given the limited potency of known fusion machinery-directed monoclonal antibodies^18,19,21,22,24^, relative to S_1_-targeting antibodies, the plasma neutralizing activity induced by SARS-CoV-2 S_2_ was weaker than that of prefusion S 2P although its breadth was much less affected by antigenic changes. In vivo evaluation of vaccine efficacy suggests that SARS-CoV-2 S_2_ protected mice comparably to prefusion S 2P, based on weight loss and viral replication in the upper and lower airways, upon challenge with the immune evasive SARS-CoV-2 XBB.1.5 variant.

We note that SARS-CoV-2 S_2_ subunit vaccination protected mice that did and those that did not have detectable serum neutralizing activity against SARS-CoV-2 XBB.1.5, as observed for stabilized MERS-CoV stems upon MERS-CoV challenge^45^. Although the immunological mechanisms underlying the observed protection remain to be defined, we postulate that weakly or non-neutralizing antibodies participated in protection through Fc-mediated effector functions, as described for the S2P6 stem-helix antibody^18^ and for S-elicited fusion machinery-directed polyclonal antibodies upon mismatched sarbecovirus challenge^46^. Exposure to SARS-CoV-2 S has also been shown to induce robust cross-reactive T cell responses which participate in protection upon exposure^47–49^ and could have played a role here as well. Future studies will decipher the contribution of these distinct branches of the adaptive immune response to the protection observed.

The broadly generalizable prefusion-stabilization strategy described here provides a robust platform for elicitation of fusion machinery-directed antibody responses and the designed antigens will enable studies of S_2_ subunit-directed immune responses. Future engineering efforts may further improve the immunogenicity of this vaccine candidate though (i) multivalent display at the surface of a nanoparticle, as exemplified for the SKYcovione SARS-CoV-2 RBD vaccine^50–52^; (ii) mRNA-launching of a membrane-anchored SARS-CoV-2 S_2_ subunit vaccine, a vaccine platform which yielded multiple efficacious SARS-CoV-2 S 2P vaccines^53^; or (iii) a recently proposed targeted deglycosylation approach of the SARS-CoV-2 S_2_ subunit to enhance neutralizing antibody titers^54^.

## Acknowledgements

This study was supported by the National Institute of Allergy and Infectious Diseases (P01AI167966 to R.B., N.P.K and D.V., DP1AI158186 and 75N93022C00036 to D.V.), a Pew Biomedical Scholars Award (D.V.), an Investigators in the Pathogenesis of Infectious Disease Awards from the Burroughs Wellcome Fund (D.V.), a Dale F. Frey Award for Breakthrough Scientists from the Damon Runyon Cancer Research Foundation (T.N.S.), the University of Washington Arnold and Mabel Beckman cryoEM center and the National Institute of Health grant S10OD032290 (to D.V.). D.V. is an Investigator of the Howard Hughes Medical Institute and the Hans Neurath Endowed Chair in Biochemistry at the University of Washington.

## Author Contributions

J.L and D.V. designed the study and the experiments; J.L and C.S. recombinantly expressed and purified glycoproteins and performed binding assays. J.L carried out pseudovirus entry assays. J.L., D.A. and Y.J.P. carried out cryoEM specimen preparation, data collection and processing. J.L. and D.V. built and refined atomic models. E.M.L. and C.T. performed mouse immunizations and blood draws. A.S. and J.M.P. carried out mice viral challenge and analysis. D.C. contributed unique reagents. J.L. and D.V. analyzed the data and wrote the manuscript with input from all authors; R.B., N.P.K., and D.V. supervised the project.

## Competing Interests

N.P.K. and D.V. are named as inventors on patents for coronavirus nanoparticle vaccines filed by the University of Washington. N.P.K. is a co-founder, shareholder, paid consultant,and chair of the scientific advisory board of Icosavax, Inc. and has received an unrelated sponsored research agreement from Pfizer. D.C. is an employee of Vir Biotechnology and may hold shares in Vir Biotechnology.

---

**Supplementary Table 1.** SARS-CoV-2 S_2_ prefusion design details with range and mutations.

| Design | Residue range | HexaPro | VFLIP inter-protomer disulfide bonds | Intra-protomer disulfide bonds |
| --- | --- | --- | --- | --- |
| **C-44** | 686-1208 | A892P, A899P, A942P, V987P | Y707C-T883C | F970C-G999C |
| Design | Residue range | Additional mutations to the C-44 background |  |  |
| **E-31** | 686-1208 | T961F |  |  |
| **E-32** | 686-1208 | D994E |  |  |
| **E-33** | 686-1208 | Q1005R |  |  |
| **E-60** | 686-1208 | F888P, A893P, N907E, T961F, D994Q, T998Q, Q1010M, Q1011M, I1018Y, N1023M |  |  |
| **E-69** | 686-1208 | N907E, T961F, D994Q, Q1011M, I1018Y |  |  |
| **F-53** | 701-1208 | N907E, T961F, D994Q, Q1011M, I1018Y |  |  |
| **SARS1 S<sub>2</sub>** | 683-1190 | N889E, T943F, D976Q, Q993M, I1000Y |  |  |
| **PRD-0038 S<sub>2</sub>** | 684-1191 | N890E, V940Q, T944F, D977Q, Q994M, I1001Y |  |  |

**Supplementary Table 2.** CryoEM data collection and refinement statistics.

|  | E-31 | E-60 | E-69 |
| --- | --- | --- | --- |
| Data collection and processing |  |  |  |
| Magnification | 105,000 | 105,000 | 105,000 |
| Voltage (kV) | 300 | 300 | 300 |
| Electron exposure (e<sup>-</sup>/Å<sup>2</sup>) | 63 | 63 | 63 |
| Defocus range (μm) | -0.2 - -7.0 | -0.2 - -3.0 | -0.1 - -3.27 |
| Pixel size (Å) | 1 | 1 | 0.843 |
| Symmetry imposed | C3 | C3 | C3 |
| Final particle images (no.) | 671,707 | 144,044 | 319,001 |
| Map resolution (Å) | 2.7 | 3.5 | 3.0 |
| FSC threshold | 0.143 | 0.143 | 0.143 |
| Map sharpening *B*factor (Å<sup>2</sup>) | -119.4 | -143.9 | -127.6 |
| Validation |  |  |  |
| MolProbity score | 1.14 | 1.11 | 1.0 |
| Clashscore | 1.29 | 1.47 | 1.33 |
| Poor rotamers (%) | 0 | 0.53 | 0.26 |
| Ramachandran plot |  |  |  |
| Favored (%) | 95.89 | 96.55 | 97.28 |
| Allowed (%) | 3.65 | 3.45 | 2.27 |
| Disallowed (%) | 0.46 | 0 | 0.45 |

**Supplementary Figure 1.**
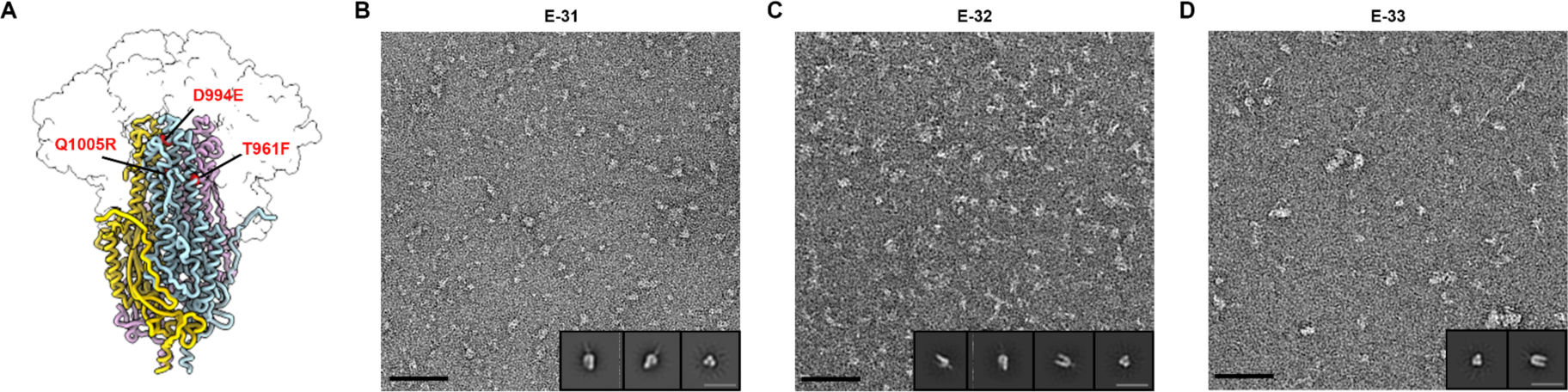
Ultrastructural characterization of designed SARS-CoV-2 S_2_ prefusion immunogens with single mutations. **A,** Ribbon diagram of prefusion SARS-CoV-2 S (PDB 6VXX^34^) highlighting all three positions shown in red (T961F, D994E, Q1005R) that were individually mutated to attempt to stabilize the metastable fusion machinery in the prefusion conformation. The S_1_ subunit is shown as a transparent surface and glycans are omitted for clarity.. **B-D,** EM analysis of negatively stained E-31 (T961F) (B), E-32 (D994E) (C), and E-33 (Q1005R) (D). Insets: 2D class averages showing compact and splayed open prefusion S_2_ trimers. The scale bar represents 50 nm (black) or 200 Å (insets, gray).

**Supplementary Figure 2.**
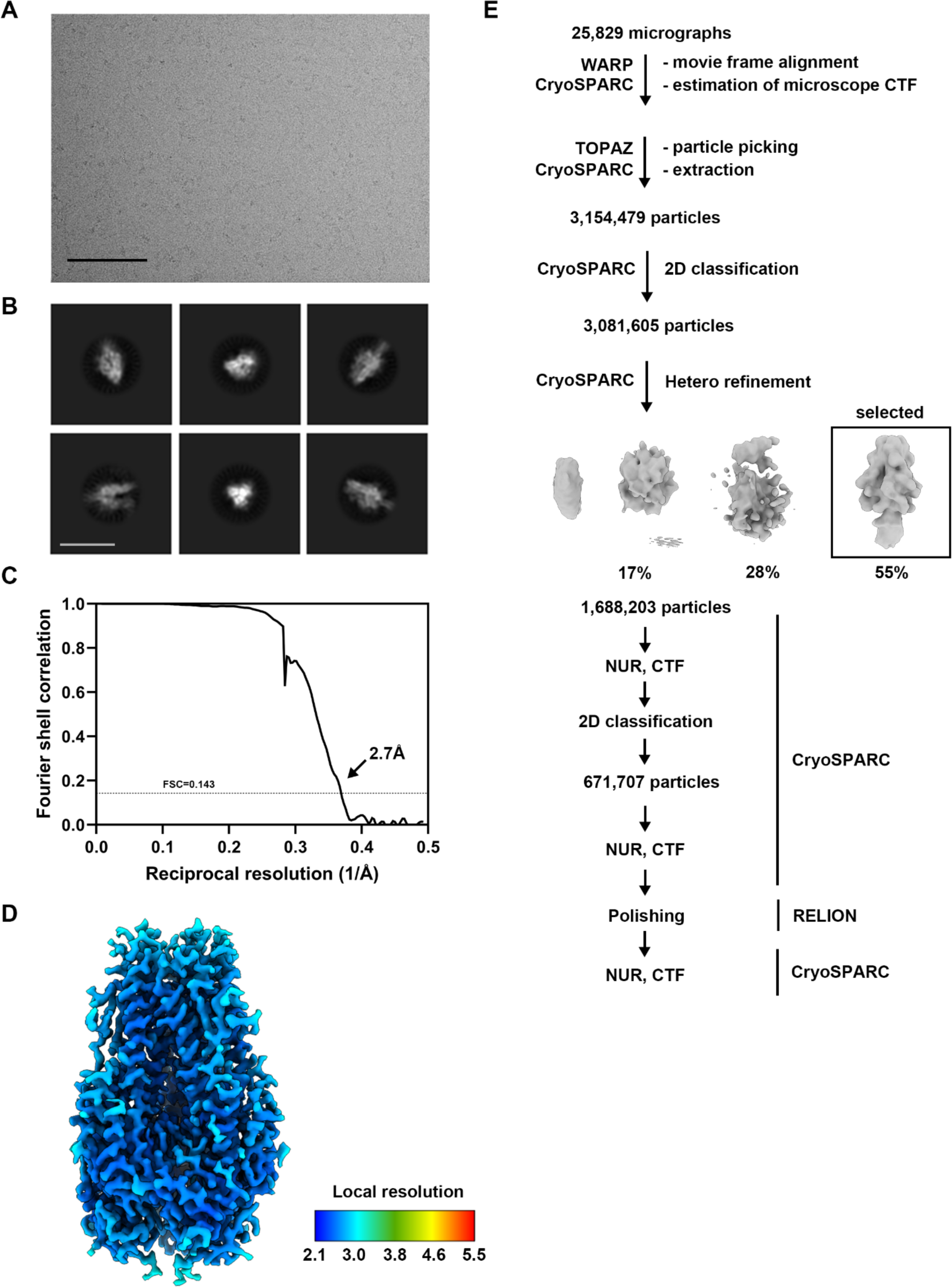
CryoEM data collection and refinement of SARS-CoV-2 S_2_ E-31. **A, B,** Representative electron micrograph (A) and 2D class averages (B) of SARS-CoV-2 S_2_ E-31 embedded in vitreous ice. The scale bar represents 100 nm (A) or 160Å (B). **C,** Gold-standard Fourier shell correlation curve for the cryoEM reconstruction. The 0.143 cutoff is indicated with a gray dashed line. **D,** SARS-CoV-2 S_2_ E-31 cryoEM map colored by local resolution as determined using cryoSPARC. **E,** Data processing flowchart. NUR, CTF: non-uniform refinement with per-particle defocus refinement.

**Supplementary Figure 3.**
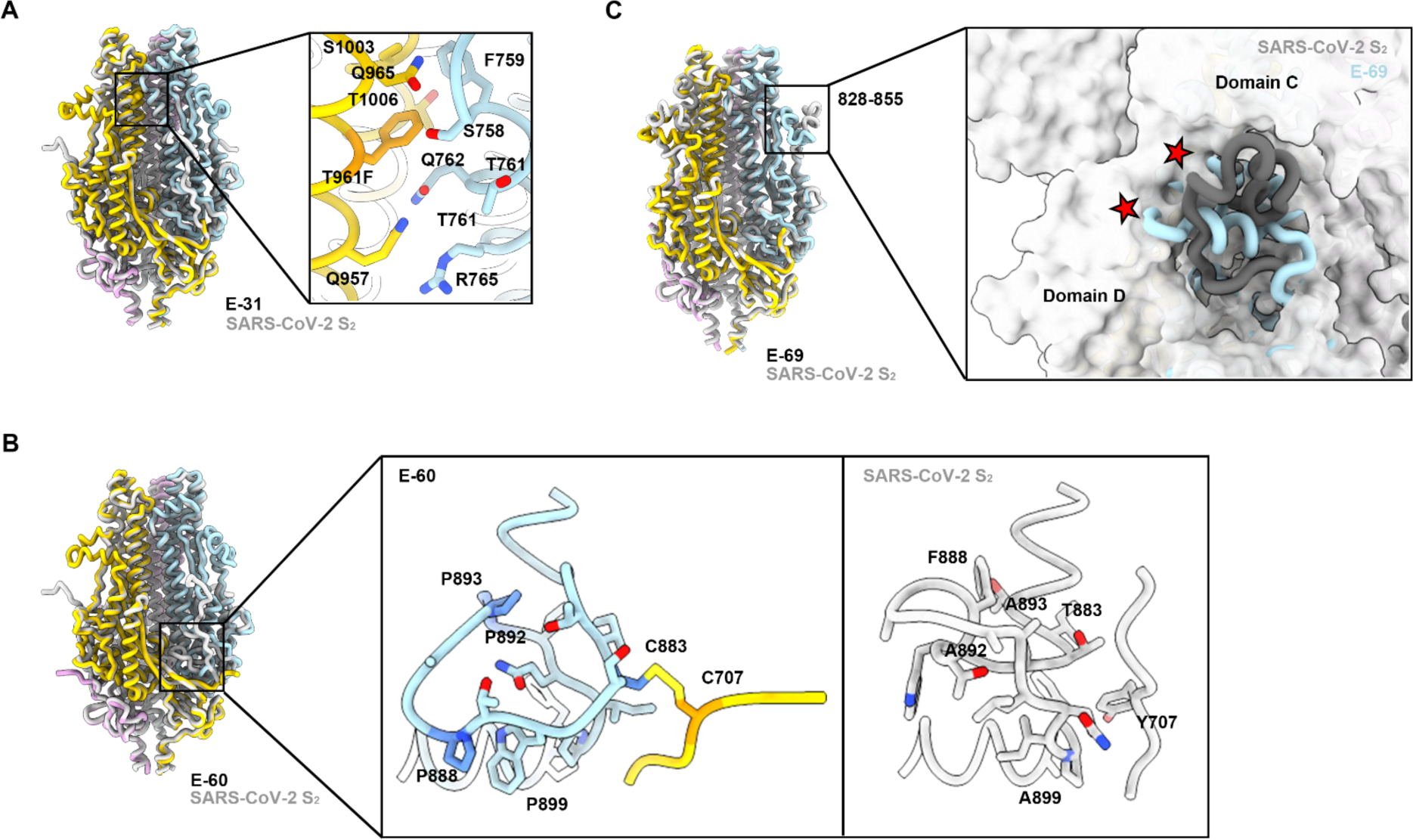
Structural details of prefusion stabilized S_2_ subunit designs. **A,** E-31 superimposed to SARS-CoV-2 S_2_ (6VXX, gray). Zoomed in view of T961F mutation and proximal residues (inset). B, E-60 superimposed to SARS-CoV-2 S_2_ (6VXX, gray). Zoomed in view of residues 875-906 of E-60 (inset, left) and SARS2 S_2_ (6VXX, gray) (inset, right). Mutated residues are shown in blue and orange. **C**, E-69 superimposed to SARS-CoV-2 S (PDB 6XR8). Steric clash shown with red stars.

**Supplementary Figure 4.**
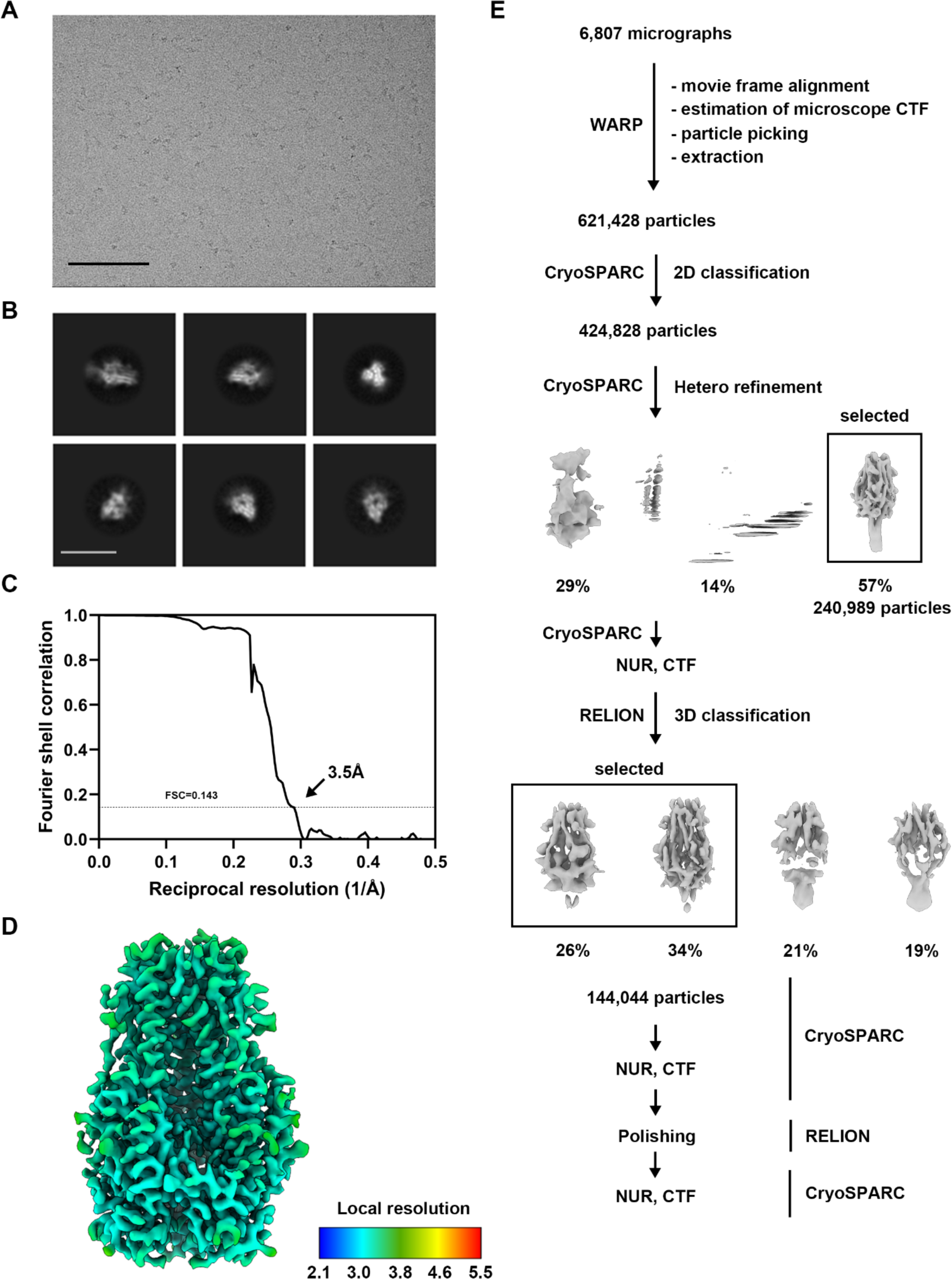
CryoEM data collection and refinement of SARS-CoV-2 S_2_ E-60. **A, B,** Representative electron micrograph (A) and 2D class averages (B) of SARS-CoV-2 S_2_ E-60 embedded in vitreous ice. The scale bar represents 100 nm (A) or 160Å (B). **C,** Gold-standard Fourier shell correlation curve for the cryoEM reconstruction. The 0.143 cutoff is indicated with a gray dashed line. **D,** SARS-CoV-2 S_2_ E-60 cryoEM map colored by local resolution as determined using cryoSPARC. **E,** Data processing flowchart. NUR, CTF: non-uniform refinement with per-particle defocus refinement.

**Supplementary Figure 5.**
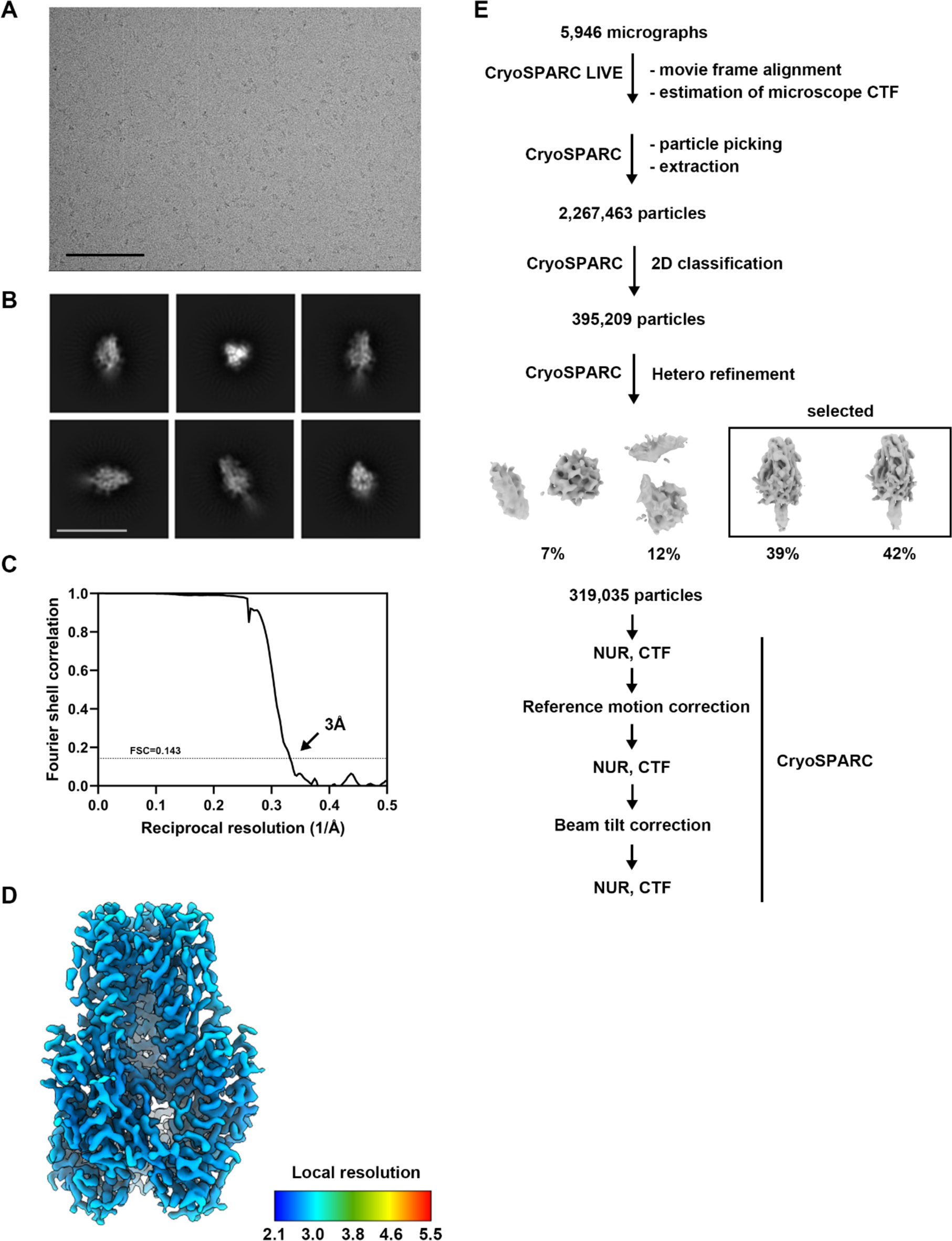
CryoEM data collection and refinement of SARS-CoV-2 S_2_ E-69. **A, B,** Representative electron micrograph (A) and 2D class averages (B) of SARS-CoV-2 S_2_ (E-69) embedded in vitreous ice. The scale bar represents 100 nm (A) or 200Å (B). **C,** Gold-standard Fourier shell correlation curve for the cryoEM reconstruction. The 0.143 cutoff is indicated with a gray dashed line. **D,** SARS-CoV-2 S_2_ (E-69) colored by local resolution as determined using cryoSPARC. **E,** Data processing flowchart. NUR, CTF: non-uniform refinement with per-particle defocus refinement.

**Supplementary Figure 6.**
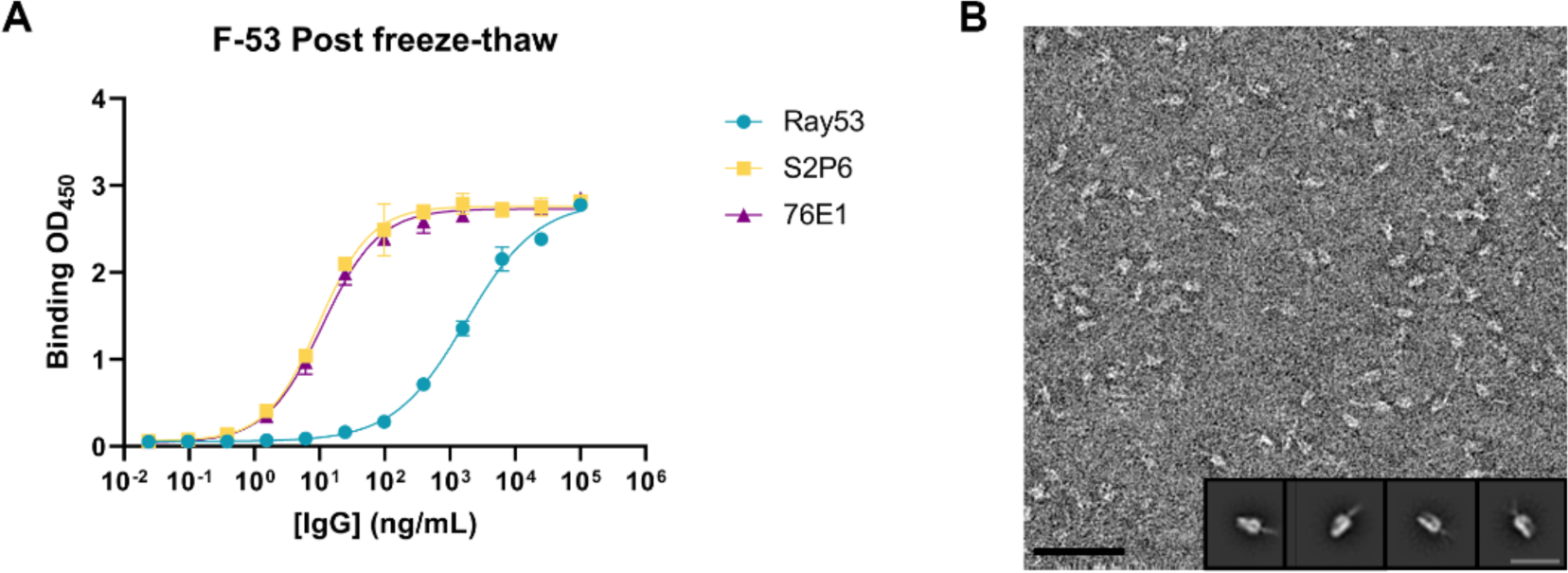
Ultrastructural characterization of SARS-CoV-2 S_2_ prefusion design F-53. **A,** Evaluation of binding of a panel of monoclonal antibodies to SARS-CoV-2 S_2_ E-69 by ELISA. **B,** EM analysis of negatively stained purified F-53. Insets: 2D class averages showing compact/splayed open prefusion S_2_ trimers. The scale bar represents 50 nm (black) or 200 Å (insets, gray).

**Supplementary Figure 7.**
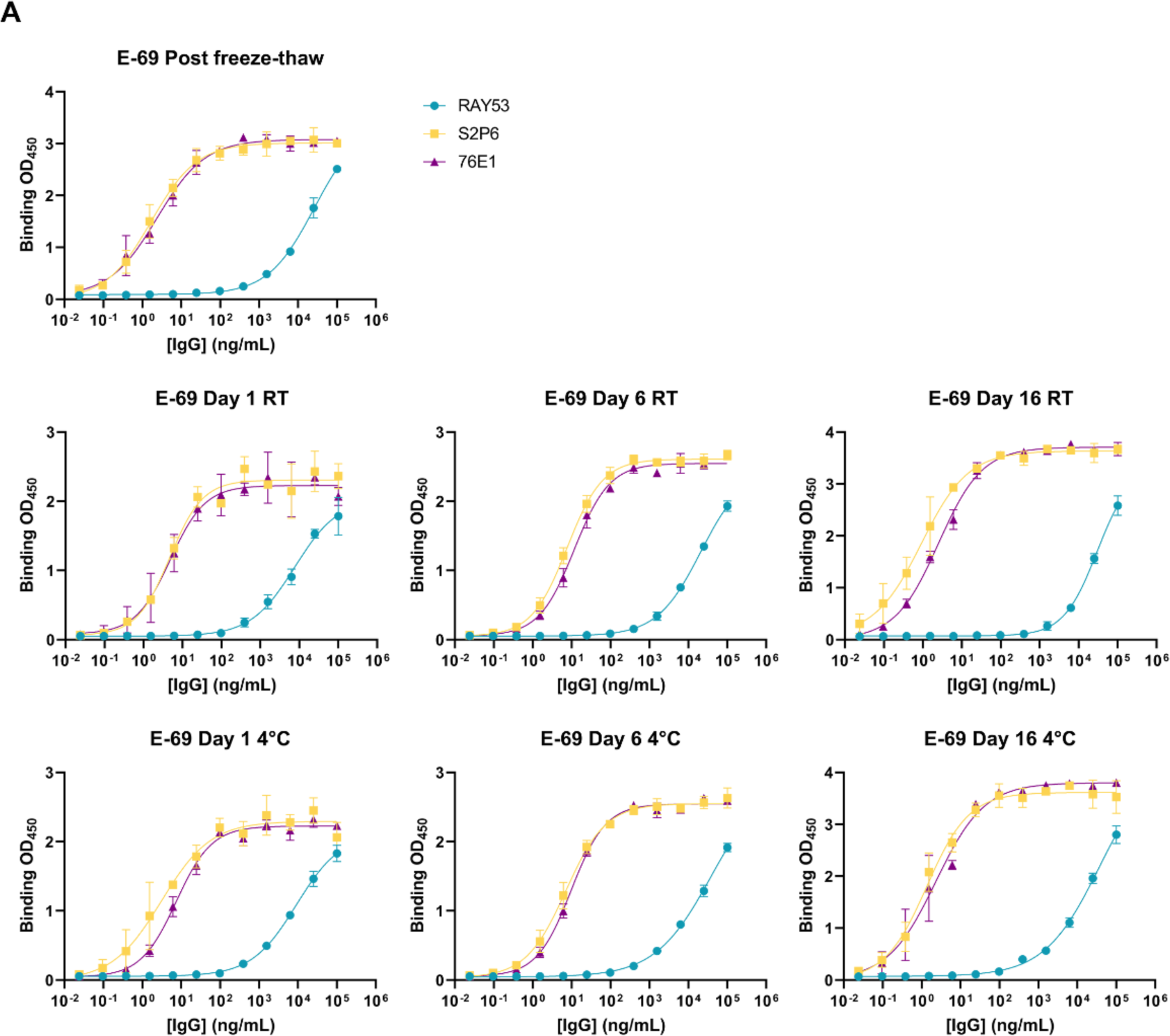
Retention of antigenicity of SARS-CoV-2 S_2_ E-69. **A,** Evaluation of binding of a panel of monoclonal antibodies to SARS-CoV-2 S_2_ E-69 under various storage conditions measured by ELISA.

**Supplementary Figure 8.**
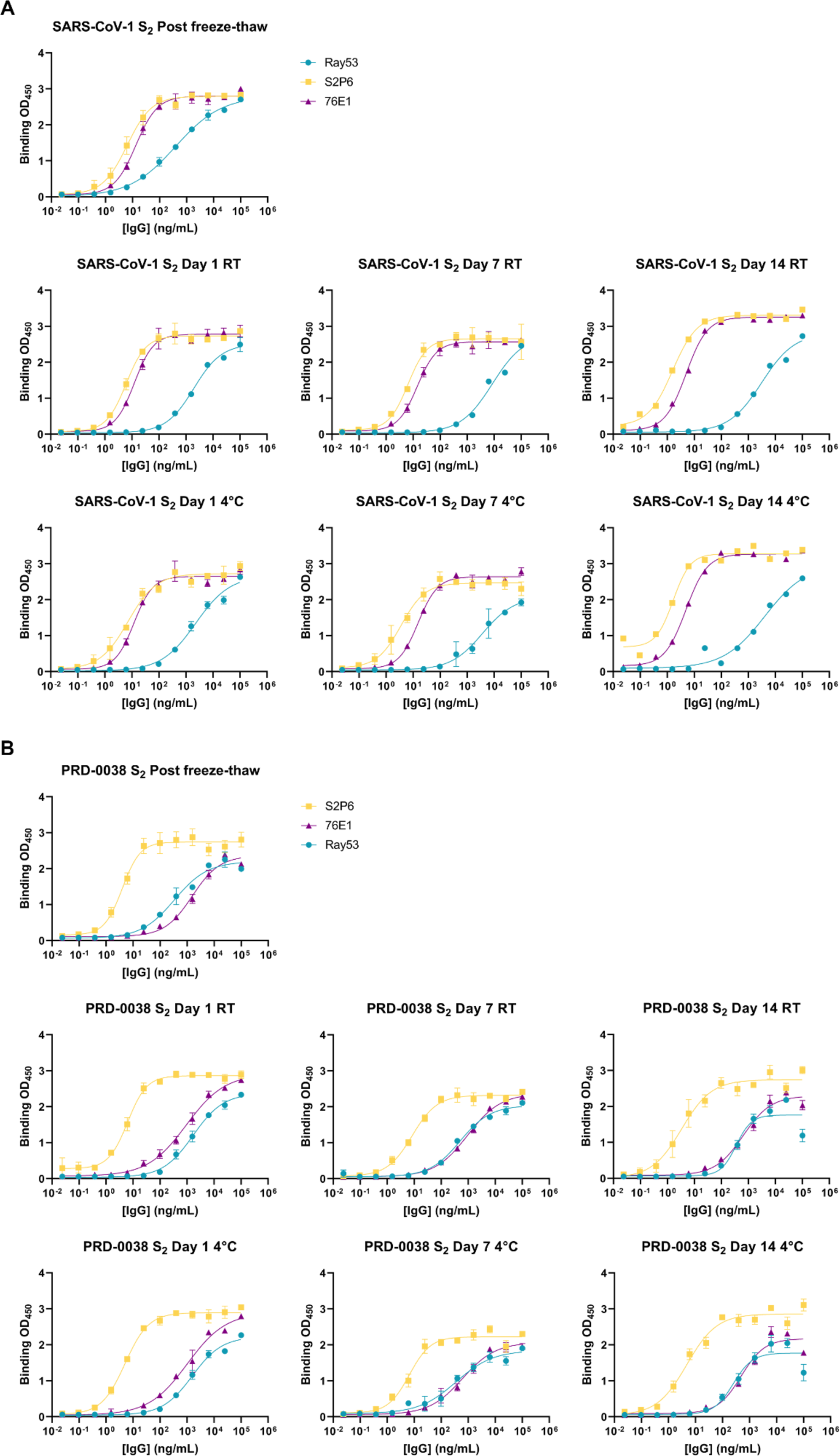
Retention of antigenicity of SARS-CoV-2 S_2_ E-69. **A,B,** Evaluation of binding of a panel of monoclonal antibodies to SARS-CoV-1 S_2_ (A) and PRD-0038 S_2_ (B) under various storage conditions measured by ELISA.

**Supplementary Figure 9.**
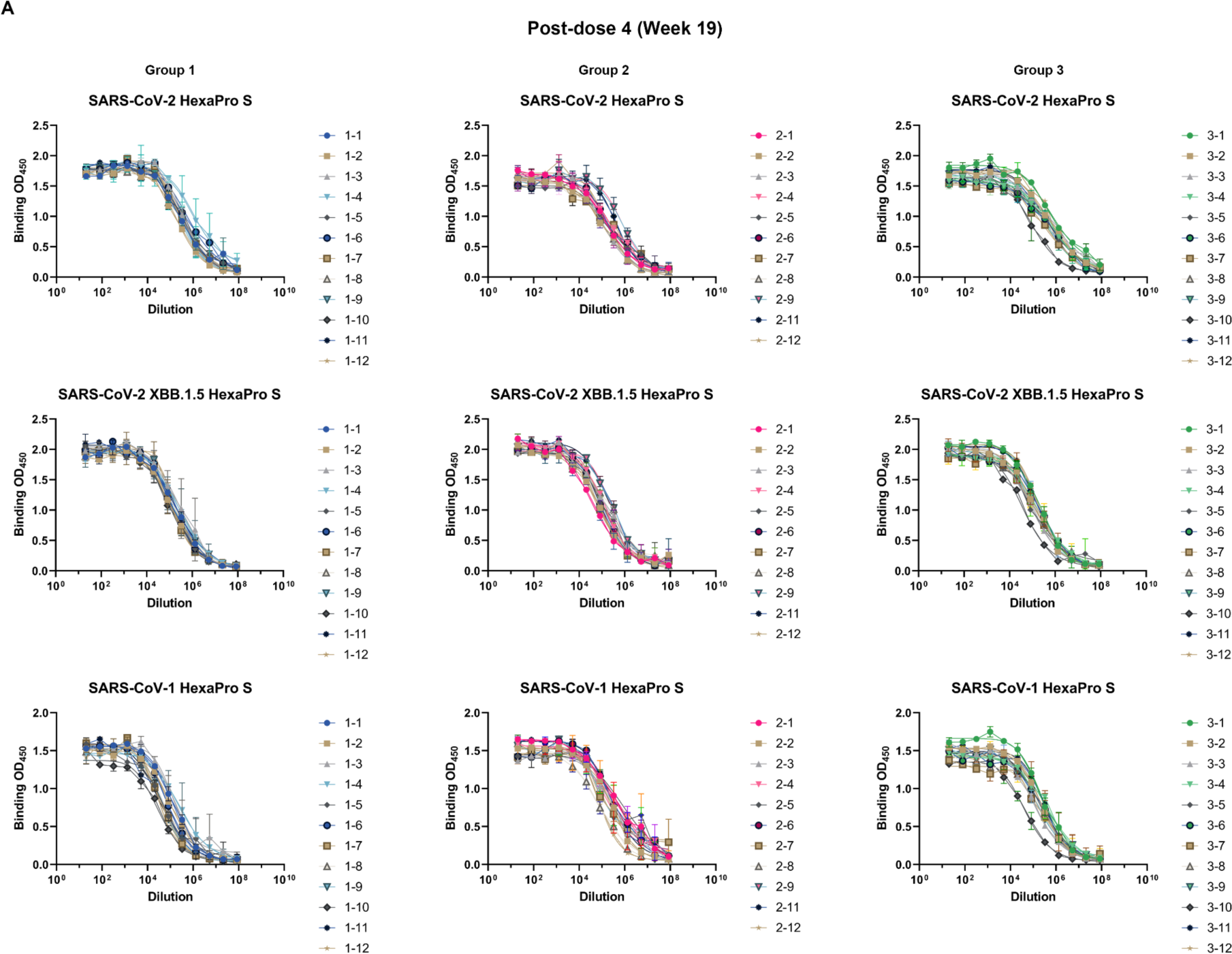
Analysis of vaccine-elicited serum antibody binding titers against various S trimers by ELISA. **A,** Representative dose-response curves of serum antibody binding to SARS-CoV-2 Hexapro S, XBB.1.5 Hexapro S, and SARS-CoV-1 Hexapro S using sera obtained 2 weeks post dose 4. Each dot represents two technical replicates. The assay has been repeated twice and the representative graph is shown.

**Supplementary Figure 10.**
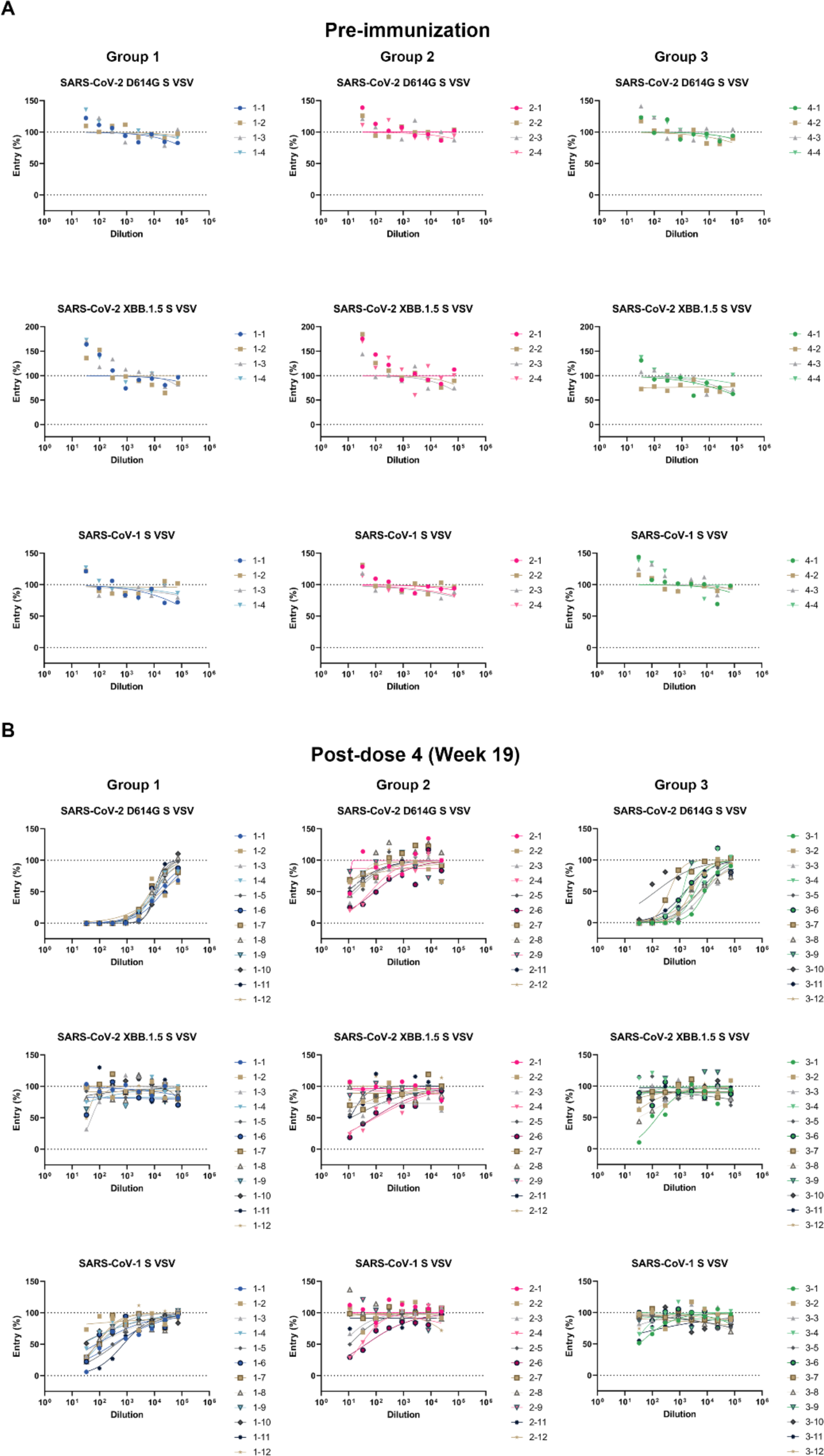
Analysis of vaccine-elicited serum neutralizing antibody titers. **A,B,** Dose-response curves of serum neutralizing antibody titers against the SARS-CoV-2 Wu/G614, XBB.1.5 and SARS-CoV-1 S VSV pseudotypes using sera obtained prior to immunization (A) and two weeks post dose 4 (B), as indicated by the color key. Representative curves from three biological replicates are shown.

## METHODS

### Cell lines

Cell lines used in this study were obtained from HEK293T (ATCC, CRL-11268), Expi293F (Thermo Fisher Scientific, A145277) and VeroE6-TMPRSS2 (JCRB1819). Cells were cultured in 10% FBS, 1% penicillin-streptomycin, 2% Geneticin (applicable for Vero cells, only) DMEM at 37℃, 5% CO_2_. None of the cell lines were authenticated or tested for mycoplasma contamination.

### Production of recombinant S_2_ antigen proteins

Each S_2_ construct was produced in Expi293F cells (ThermoFisher Scientific) and cultured at 37°C in a humidified 8% CO_2_ incubator with constant rotation at 130 RPM using Expi293 Expression Medium (ThermoFisher Scientific). DNA transfections were conducted using the ExpiFectamine 293 Transfection Kit (ThermoFisher Scientific) protocols and materials and cultivated for five days before harvest. Cell culture supernatants were clarified by centrifugation and proteins were harvested using HisTrap™ High Performance Ni Sepharose columns (Cytiva). Proteins were washed using 10-15 CVs of buffer containing 25 mM Tris, 150 mM NaCl, 20 mM Imidazole pH 8.0 followed by elution with 10-15 CVs of buffer containing 25 mM Tris, 150 mM NaCl, 300 mM Imidazole pH 8.0. Eluates were buffer exchanged and concentrated into 20 mM Tris, 150 mM NaCl, pH 8.0 using Amicon Ultra-15 Centrifugal Filter Unit (10 kDa) (Millipore). Gel filtration was performed to remove unfolded or aggregated protein thus samples were each run through a Superose-6 Increase 10/300 GL column (Cytiva)) equilibrated in 20 mM Tris, 150 mM NaCl, pH 8.0. Main peaks were collected and protein was snap frozen and stored at -80 °C with some set aside for stability tests. Purified proteins for immunogenicity study were tested for endotoxin levels using Limulus Amebocyte Lysate (LAL) cartridges (Charles River PTS201F).

### Production of recombinant SARS-CoV-2 HexaPro S and SARS-CoV-1 HexaPro S

The SARS-CoV-2 HexaPro S glycoprotein ectodomain construct comprises residues 1-1208 with the native signal peptide, the HexaPro prefusion stabilizing mutations (F817P, A892P, A899P, A942P, K986P, V987P), abrogation of the furin cleavage site (residues 682-685, GSAS) and a C-terminal short linker (GSG), followed by a foldon, HRV 3C site (LEVLFQGP), a short linker (GSG), an avi tag, a short linker (GSG), an 8x his tag in a pcDNA3.1(-) plasmid. The SARS-CoV-1 HexaPro S glycoprotein was expressed as previously described. Expi293F cells were grown at 37℃ with 8% CO_2_ and DNA transfections were conducted with the ExpiFectamine 293 Transfection Kit (Thermo Fisher Scientific). Cell culture supernatants were harvested four days post-transfection and proteins were purified using HisTrap™ High Performance column (Cytiva). Proteins were first washed with 10-15 column volumes of a buffer containing 25 mM sodium phosphate, 300 mM NaCl, 20 mM imidazole, pH 8.0, followed by elution with 10-15 column volumes using 300 mM imidazole, pH 8.0. Eluted proteins were concentrated and buffer exchanged into 1x TBS (20 mM Tris, 150 mM NaCl, pH 8.0) using Amicon Ultra-15 Centrifugal Filter Unit (100 kDa) (Millipore). Purified proteins were snap frozen and stored at -80°C.

### Monoclonal antibody ELISAs

For monoclonal antibody ELISAs, 30 μl of the proteins at 3 μg/mL were plated onto 384-well Nunc Maxisorp plate (ThermoFisher, 464718) in 1x TBS and incubated 1h at 37°C followed by slap drying and blocking with 80 μL of Casein for 1 h at 37°C. After incubation, plates were slap dried and 1:4 serial dilutions of the corresponding mAbs starting from 0.1 mg/ml were made in 30 μl TBST, added to the plate and incubated at 37°C for 1 h. Plates were washed 4x in TBST and 30 μl of 1:5,000 Goat anti-Human IgG Fc Secondary Antibody, HRP (Thermo Fisher, A18817) were added to each well and incubated at 37°C. After 1 h, plates were washed 4x in TBST and 30 μl of TMB (SeraCare) was added to every well for 2 min at room temperature. Reactions were quenched with the addition of 30 μl of 1N HCl. Plates were immediately read at 450 nm on a BioTek Neo2 plate reader and data plotted and fit in Prism 9 (GraphPad) using nonlinear regression sigmoidal, 4PL, X is the concentration to determine EC_50_ values from curve fits.

### Production of VSV pseudoviruses

SARS-CoV-2 D614G S, XBB.1.5 S, and SARS-CoV-1 S VSV pseudoviruses were produced using HEK293T cells seeded on BioCoat Cell Culture Dish : poly-D-Lysine 100 mm (Corning). Cells were transfected with respective S constructs using Lipofectamine 2000 (Life Technologies) in Opti-MEM transfection medium. After 5h of incubation at 37 °C with 5% CO2, cells were supplemented with DMEM containing 10% of FBS. On the next day, cells were infected with VSV (G*ΔG-luciferase) for 2h, followed by five time wash with DMEM medium before addition of anti-VSV G antibody (I1-mouse hybridoma supernatant diluted 1:40, ATCC CRL-2700) and medium. After 18-24 h of incubation at 37 °C with 5% CO_2_, pseudoviruses were collected and cell debris removed by centrifugation at 3,000xg for 10 min. Pseudoviruses were further filtered using a 0.45 µm syringe filter and concentrated 10x prior to storage at -80°C.

### Serological ELISAs

For serological ELISAs, 30 µL of assorted proteins (SARS-CoV-2 HexaPro S, SARS-CoV-2 XBB.1.5 HexaPro S (AcroBiosystems, SPN-C524i) and SARS-CoV-1 HexaPro S) at 3 µg/mL were placed into 384-well Nunc Maxisorp plates (ThermoFisher, 464718) in 1x TBS and incubated for 1 hour at 37°C followed by slap drying and blocking with 80 µL of Casein for 1 hour at 37°C. Afterwards, plates were once again slap dried and a 1:4 serial dilution of our immunized mouse sera was performed starting from 1:20 dilution in 30 µL of TBST and incubated at 37°C for 1 hour. Plates were then washed 4x in TBST and 30 µL of 1:5,000 Goat anti-mouse IgG (H+L) Secondary Antibody HRP (ThermoFisher 62-6520) were added to each well and incubated at 37°C for 1 hour. Plates were then washed 4x in TBST and 30 uL of TMB (SeraCare) was added to each well and allowed to sit for 2 minutes at room temperature. TMB reactions were quenched with 30 µL of 1N HCl and immediately read at 450 nm on a BioTek Neo2 plate reader and data plotted and fit in Prism 10 (Graphpad) using nonlinear regression sigmoidal, 4PL, X is the concentration to determine ED_50_ values from curve fits.

### Negative stain electron microscopy preparation, data collection, and data processing

Carbon copper formvar grids (Ted Pella 01754-F) were glow discharged using a Gloqube Plus (Quorum) at 20 mA for 30 seconds promptly followed by the addition of 3 µL of a S_2_ pre-fusion constructs diluted to a concentration of 0.01 mg/mL. After 1 minute the protein was aspirated using filter paper and 3 µL of 2% uranyl formate was applied and quickly removed for washing. Another 3 µL of uranyl formate was added to the grid and left to stain for 30 seconds before drying with filter paper and left to further air dry before imaging. Automated data collection was carried out using Leginon at a nominal magnification of 67,000 with a pixel size of 1.6 Å. Each micrograph was acquired for 500-900 ms. Negative stain data was processed using CryoSPARC. Automatic particle picking and extraction were performed using CryoSPARC for each data set. Particle images were extracted with a box size of 256 pixels with a pixel size of 1.6Å and binned to 128 pixels for subsequent 2D classifications.

### Cryo-EM sample preparation and data collection

The E-31 cryo-EM dataset was collected at three different times and combined to be processed together. 3 µL of sample was added to a glow discharged (120s at 20mA) UltraAuFoil R2/2:Au200 grid prior to plunge freezing using a vitrobot MarkIV (ThermoFisher Scientific) with a blot force of 0 and 6.5 sec blot time at 100 % humidity and 22°C. For E-60, 3 µL of sample was added to a glow discharged (120s at 20mA) UltraAuFoil R2/2:Au200 grid prior to plunge freezing using a vitrobot MarkIV (ThermoFisher Scientific) with a blot force of 0, 5.5 sec blot time, and 10s wait time at 100 % humidity and 22°C. For E-69, 3 µL of sample was added to a glow discharged (120s at 20mA) UltraAuFoil R2/2:Au200 grid prior to plunge freezing using a vitrobot MarkIV (ThermoFisher Scientific) with a blot force of 0, 6 sec blot time, and 10s wait time at 100 % humidity and 22°C.

Data were acquired using an FEI Titan Krios transmission electron microscope operated at 300 kV and equipped with a Gatan K3 direct detector and Gatan Quantum GIF energy filter, operated in zero-loss mode with a slit width of 20 eV. Automated data collection was carried out using Leginon^55^ at a nominal magnification of 105,000x with a pixel size of 0.843 Å. The dose rate was adjusted to 15 counts/pixel/s, and each movie was acquired in counting mode fractionated in 75 frames of 40 ms. A total of 25,829 and 6,807 micrographs were collected for E-31 and E-60 datasets,respectively. Stage was tilted 0, 30, and 45 degrees for E-31 and 20 degrees for the E-60 collection. A total of 5,946 micrographs were collected for E-69 at 0 and 30 degrees tilted stage.

### Cryo-EM data processing, model building and refinement

For the E-31 structure, motion correction and contrast-transfer function (CTF) parameter estimation were performed using Warp^56^ and cryoSPARC, respectively. Automatic particle picking was performed using TOPAZ^57^ and particle images were extracted with a box size of 208 pixels with a pixel size of 1.686Å. After 2D classification and hetero-refinement using cryoSPARC^58^, 1,688,203 particles were selected for cryoSPARC non-uniform refinement^59^ with C3 symmetry. Particles were further subjected to another round of 2D classification followed by Bayesian polishing^60^ in Relion. Finally, another round of cryoSPARC non-uniform refinement with C3 symmetry and per-particle defocus refinement was carried out using the polished particles.

For the E-60 structure, motion correction, CTF estimation, automatic particle picking, and extraction were performed using Warp^56^. Particle images were extracted with a box size of 208 pixels and a pixel size of 1.686Å. After 2D classification and hetero-refinement in cryoSPARC, 240,989 particles were selected. These particles were subjected to two rounds of 3D classification with 50 iterations each (angular sampling 7.5° for 25 iterations and 1.8° with local search for 25 iterations) using Relion^61–63^. 3D refinements were carried out using non-uniform refinement along with per-particle defocus refinement in cryoSPARC followed by Bayesian polishing^60^ in Relion. Finally, another round of cryoSPARC non-uniform refinement with C3 symmetry and per-particle defocus refinement was carried out using the polished particles.

For the E-69 structure, motion correction, CTF estimation, automatic particle picking, and extraction were performed using cryoSPARC LIVE and cryoSPARC. Particle images were extracted with a box size of 208 pixels and a pixel size of 1.686Å. After 2D classification and hetero-refinement in cryoSPARC, 319,035 particles were selected and subjected to reference motion correction and beam tilt correction followed by a final non-uniform refinement with C3 symmetry using cryoSPARC.

Local resolution estimation, filtering, and sharpening were carried out using CryoSPARC. Reported resolutions are based on the gold-standard Fourier shell correlation (FSC) of 0.143 criterion and Fourier shell correlation curves were corrected for the effects of soft masking by high-resolution noise substitution^64,65^. UCSF Chimera^66^, UCSF ChimeraX^67^, and Coot^68^ were used to fit atomic models into the cryoEM maps. The built models were refined and relaxed using Rosetta^69,70^ using sharpened and unsharpened maps and validated using Phenix^71^, Molprobity^72^ and Privateer^73^.

### Immunogenicity

Female BALB/c mice were purchased from Envigo (order code 047) at 7 weeks of age and were maintained in a specific pathogen-free facility within the Department of Comparative Medicine at the University of Washington, Seattle, accredited by the Association for Assessment and Accreditation of Laboratory Animal Care (AAALAC). Animal experiments were conducted in accordance with the University of Washington’s Institutional Animal Care and Use Committee.

Prior to each immunization, immunogens (low endotoxin immunogen) were diluted to 20 µg/mL (S2P) or 100 µg/mL (S_2_) in 1x PBS (1.5 mM Potassium Phosphate monobasic, 155mM NaCl, 2.7mM Sodium Phosphate diabasic, pH 7.4) (ThermoFisher) and mixed with 1:1 vol/vol AddaVax (InvivoGen vac-adx-10) to reach a final dose of 1 µg (S2P) or 5µg (S_2_) of immunogen per injection. At 8 weeks of age, 12 mice per group were injected subcutaneously in the inguinal region with 100 uL of immunogen at weeks 0, 3, 10, and 17. Group 1 received four doses of 1 µg S2P. Group 2 received four doses of 5 µg S_2_. Group 3 received two doses of 1 µg S2P and boosted with two doses of 5 µg S_2._ Mice were bled via the submental route at weeks 0, 2, 5, 12, and 19. Blood was collected in serum separator tubes (BD # 365967) and rested for 30 min at room temperature for coagulation. Serum tubes were then centrifuged for 10 min at 2,000 x g and serum was collected and stored at -80°C until use.

### Neutralization assays

For SARS-CoV-2 D614G S VSV, XBB.1.5 S VSV, and SARS-CoV-1 S VSV neutralization, VeroE6-TMPRSS2 cells in DMEM supplemented with 10% FBS, 1% PenStrep, and 2% Geneticin were seeded at 40,000 cells/well into 96-well plates [3610] (Corning) and incubated overnight at 37°C. The following day, a half-area 96-well plate (Greiner) was prepared with 3-fold serial sera dilutions (starting dilutions determined for each serum and pseudovirus, 22uL per well). An equal volume of DMEM with diluted pseudoviruses was added to each well. All pseudoviruses were diluted between 1:90-1:200 to reach a target entry of ∼10^6^ RLU. The mixture was incubated at room temperature for 45-60 minutes. Media was removed from the cells and the cells were washed once with DMEM prior to the transfer of sera-pseudovirus mixture. 40 μL from each well of the half-area 96-well plate containing sera and pseudovirus were transferred to the 96-well plate seeded with cells and incubated at 37°C for 1h. After 1h, an additional 40 μL of DMEM supplemented with 20% FBS and 2% PenStrep was added to the cells. After 18–20h, 40 μL of One-Glo-EX substrate (Promega) was added to each well and incubated on a plate shaker in the dark for 5 min before reading the relative luciferase units using a BioTek Neo2 plate reader. Relative luciferase units were plotted and normalized in Prism (GraphPad): 100% neutralization being cells lacking pseudovirus and 0% neutralizing being cells containing virus but lacking sera. Prism (GraphPad) nonlinear regression with “log[inhibitor] versus normalized response with a variable slope” was used to determine ID_50_ values. 3 biological replicates were carried out for each sample-pseudovirus pair.

## Notes

### Summary of Updates

Challenge data controls were missing

